# ProkBERT PhaStyle: Accurate Phage Lifestyle Prediction with Pretrained Genomic Language Models

**DOI:** 10.1101/2024.12.08.627378

**Authors:** Judit Juhász, Bodnár Babett, János Juhász, Noémi Ligeti-Nagy, Sándor Pongor, Balázs Ligeti

**Affiliations:** Faculty of Information Technology and Bionics, Pázmány Péter Catholic University, Práter st. 50/a, 1083, Budapest, Hungary; Institute of Medical Microbiology, Semmelweis University, Nagyvárad sq. 4, 1089, Budapest, Hungary; Language Technology Research Group, HUN-REN Hungarian Research Centre for Linguistics, Benczúr st. 33, 1068, Budapest, Hungary

**Keywords:** phage lifestyle prediction, genomic language models, ProkBERT, virulent phages, transfer learning

## Abstract

**Background:** Phage lifestyle prediction, i.e. classifying phage sequences as virulent or temperate, is crucial in biomedical and ecological applications. Phage sequences from metagenome or metavirome assemblies are often fragmented, and the diversity of environmental phages is not well known. Current computational approaches often rely on database comparisons and machine learning algorithms that require significant effort and expertise to update. We propose using genomic language models for phage lifestyle classification, allowing efficient direct analysis from nucleotide sequences without the need for sophisticated preprocessing pipelines or manually curated databases.

**Methods:** We trained three genomic language models (DNABERT-2, Nucleotide Transformer, and ProkBERT) on datasets of short, fragmented sequences. These models were then compared with dedicated phage lifestyle prediction methods (PhaTYP, DeePhage, BACPHLIP) in terms of accuracy, prediction speed, and generalization capability.

**Results:** ProkBERT PhaStyle consistently outperforms existing models in various scenarios. It generalizes well for out-of-sample data, accurately classifies phages from extreme environments, and also demonstrates high inference speed. Despite having up to 20 times fewer parameters, it proved to be better performing than much larger genomic language models.

**Conclusions:** Genomic language models offer a simple and computationally efficient alternative for solving complex classification tasks, such as phage lifestyle prediction. ProkBERT PhaStyle’s simplicity, speed, and performance suggest its utility in various ecological and clinical applications.

## Introduction

Bacteriophages, the viruses targeting bacteria, exhibit distinct lifestyles: virulent phages destroy host bacteria through the lytic cycle, potentially clearing pathogenic bacterial infections quickly; temperate phages on the other hand integrate into the bacterial genome and can transfer genes, influencing bacterial evolution and pathology [1]. Understanding phage-host interactions is pivotal for advancing medical and environmental biotechnology [2, 3], such as phage therapeutic applications [4, 2, 5], and microbiome engineering [6, 7].

Despite the critical role of phages, predicting their phenotypes, particularly their lifestyles, remains less explored. Existing tools for phage lifestyle prediction can be categorized into nucleotide-based and protein-based approaches. Temperate and virulent phages possess distinct characteristics: temperate phages often encode toxins, integrase, and excisionase genes, whereas virulent phages typically encode lysis and nucleotide metabolism-related genes [8, 9]. These features form the basis for heuristic-driven phage lifestyle predictors. Protein-based tools, such as PHACTS, utilize protein information and machine learning techniques like Random Forests to predict phage lifestyles [10]. BACPHLIP [11], which uses HMM profiles to identify lysogeny-associated protein domains, and PhaTYP [12], which relies on database searches to construct protein sentences for prediction, are examples of this approach. Conversely, nucleotide-based tools like PhagePred utilize a k-mer frequency-based Markov model to calculate the similarity of query contigs to known temperate and virulent phages [13]. DeePhage, an alignment-free approach, employs a convolutional neural network to learn local patterns, enabling it to classify contigs effectively [14].

Several challenges persist in phage lifestyle prediction: (i) accurately labeling fragmented phage sequences, (ii) achieving computational efficiency, and (iii) recognizing and classifying phages that are underrepresented or absent in the training set. Phage sequences derived from metaviromic and metagenomic studies are often incomplete and fragmented, reducing the accuracy of current methods such as BACPHLIP, PHACTS, and PhagePred for shorter fragments (*<*5kb) [14, 13]. Despite advancements in viral catalogs [15, 16, 17, 18], the human virome remains under-characterized [19]. This leads to biases and computational expenses in database-based approaches such as PhaTYP and BACPHLIP, which often fail in ’out-of-sample’ scenarios, where they encounter previously unseen phages [20]. Addressing these challenges is essential for developing robust, generalizable phage lifestyle prediction models.

Transformer-based language models have demonstrated excellent generalization capabilities in natural language processing (NLP) tasks [21], making them good candidates for data-scarce scenarios common in biomedical fields. In biological sequences, transformer models such as MSA Transformer [22], ESM [23, 24], AlphaFold2 [25], and openFold [26] have been effectively applied. For nucleotide sequences, models such as DNABERT [27, 28], GENA-LM [29], Nucleotide Transformer [30] (NT), Mamba [31], HyenaDNA [32], and ProkBERT, a model pretrained on microbial sequences [33] show promise but they have not yet been employed to phage lifestyle prediction tasks.

In this work we fine-tune universal genomic language models (Nucleotide Transformer: 50-500m parameters, DNABERT-2: 117m, and ProkBERT: 21-26m parameters) and then benchmark them against the current state of the art phage lifestyle prediction tools (BACPHLIP, DeePhage, PhaTYP). The benchmarking focuses on 3 aspects, i) how well the pretrained language models can classify fragments as short as 512bp, ii) the prediction speed of each method and iii) how well the models perform on ’unseen’, out-of-the sample predictions.

We demonstrate that the task of phage lifestyle prediction can be solved efficiently and with high classification accuracy directly from nucleotide sequences using a simple pipeline. This approach is fast, requires no arbitrary thresholding, and provides improved generalization by leveraging the advantages of large genomic language model (LM)-derived sequence representations. Our results show that ProkBERT PhaStyle substantially outperforms existing models in scenarios involving unseen phage groups and fragmented sequences, highlighting its potential as a robust tool for phage lifestyle prediction in diverse ecological and clinical settings.

## Data and Methods

The efficacy and applicability of machine learning algorithms are profoundly impacted by the quality of the training dataset and the comprehensiveness of the benchmarking process used for model optimization. A recurrent challenge in bioinformatics is the application of developed computational tools to sequences that were not represented or were underrepresented in the model’s training data, potentially leading to biased predictions. To address this issue and simulate real-world application scenarios, our study specifically focuses on testing models against ’unseen’ genera, with an emphasis on *Escherichia* and extremophile groups.

### Datasets

#### Training and validation datasets

The training and validation datasets were assembled to ensure high-quality annotations and a broad representation of phage lifestyles, building on the work of Mavrich & Hatfull (2017) and the BACPHLIP study by Hockenberry and Wilke [34, 20]. These datasets include 2,114 sequences: 1,868 in the training set (non-*Escherichia* phages) and 246 in the validation set (*Escherichia* phages).

#### Test dataset 1 –– Guelin (Escherichia) collection

A collection of 96 taxonomically diverse *Escherichia* bacteriophages (Guelin collection) was used for testing the methods. This collection was constructed by Gaborieau et al. [35] for experimentally studying virulent host-phage interactions in different phage concentrations. It contains the archetypical T4 and T7 phages of *E. coli* and 94 other phages isolated from different wastewater treatments plants around Paris (France) over 10 years. Their lytic capabilities were experimentally evaluated against different members of the *Escherichia* genus. The collection contains unique members with less than 98% nucleotide identity or different host-range.

#### Test dataset 2 –– Extremophile collection

This test dataset is a curated collection of bacteriophages isolated from extreme environments (see Supplementary Table 1) – deep-sea (Mariana Trench), acidic and arsenic-rich environments. The virulent phages live in arsenic-rich microbial mats in cold climates [36]. The temperate phages infect the psychrotolerant deep-sea bacterium *Aurantimonas* and *Halomonas* [37, 38, 39]. The collection also contains a bacteriophage with hosts like *Acidithiobacillus caldus*, an extremophilic bacterium thriving in highly acidic environments (pH *<* 2) [40].

#### Application of current phage lifestyle prediction methods

DeePhage employs one-hot encoding for vectorizing individual sequences. The implementation relies on Keras and MATLAB; therefore, we encapsulated DeePhage into both Docker and Apptainer containers. DeePhage was used with the default settings and GPU support. All datasets were evaluated at both the sequence and segment levels.

PhaTYP uses a BERT architecture for classification, but it operates on protein sentences, rather than nucleotide or amino-acid sequences. It was used with multithread support and was encapsulated into an Apptainer. The segment datasets were evaluated separately from the complete sequences.

BACPHLIP was installed via pip (version 0.9.6). The ’multi fasta’ option was used for evaluating the sequence datasets. However, the segment datasets were not evaluated, as this is against the recommended usage by the authors.

All Dockerfiles, Singularity definition files are available on the GitHub page. The Docker containers are also available on Docker Hub, and the Apptainer images are shared on Zenodo.

#### Running time and inference speed estimation

The running time was measured on a set of 1,000 randomly selected sequences from the BACPHLIP dataset. We evaluated methods that support GPU inference (genomic LMs, PhaTYP, DeePhage), with each model allocated one NVIDIA Tesla A100 GPU, 8 CPU cores, and 32 GB of RAM. The batch size for evaluation was adjusted based on the input sequence length and model size, as described previously. The total elapsed time accounted for all processing stages, including initialization, model and dataset loading, parsing, preprocessing of the sequence datasets (such as segmentation and tokenization), inference, and report generation (including weighted voting). Each measurement was independently repeated three times under identical conditions, and the average running time was reported. The inference speed is defined as the rate of classification (number of classified nucleotides per second).

#### Evaluation Metrics

We used standard metrics to evaluate binary classification performance, including balanced accuracy, accuracy, sensitivity, specificity, F1 score, and MCC (Matthews Correlation Coefficient). The cross-entropy loss function was used during the training of the genomic LM. Formal definitions of these metrics can be found in the supplementary material (section: Evaluation Metrics and Definitions).

## Results and Discussion

### Data preprocessing

The results of metagenome and metavirome assemblies are often fragmented, especially for low-coverage phages. Such cases are especially relevant for less known environmental samples. To accommodate these cases, we trained the models specifically to classify fragmented contigs as well. The dataset preprocessing process is summarized on Figure 1.

**Fig. 1.**
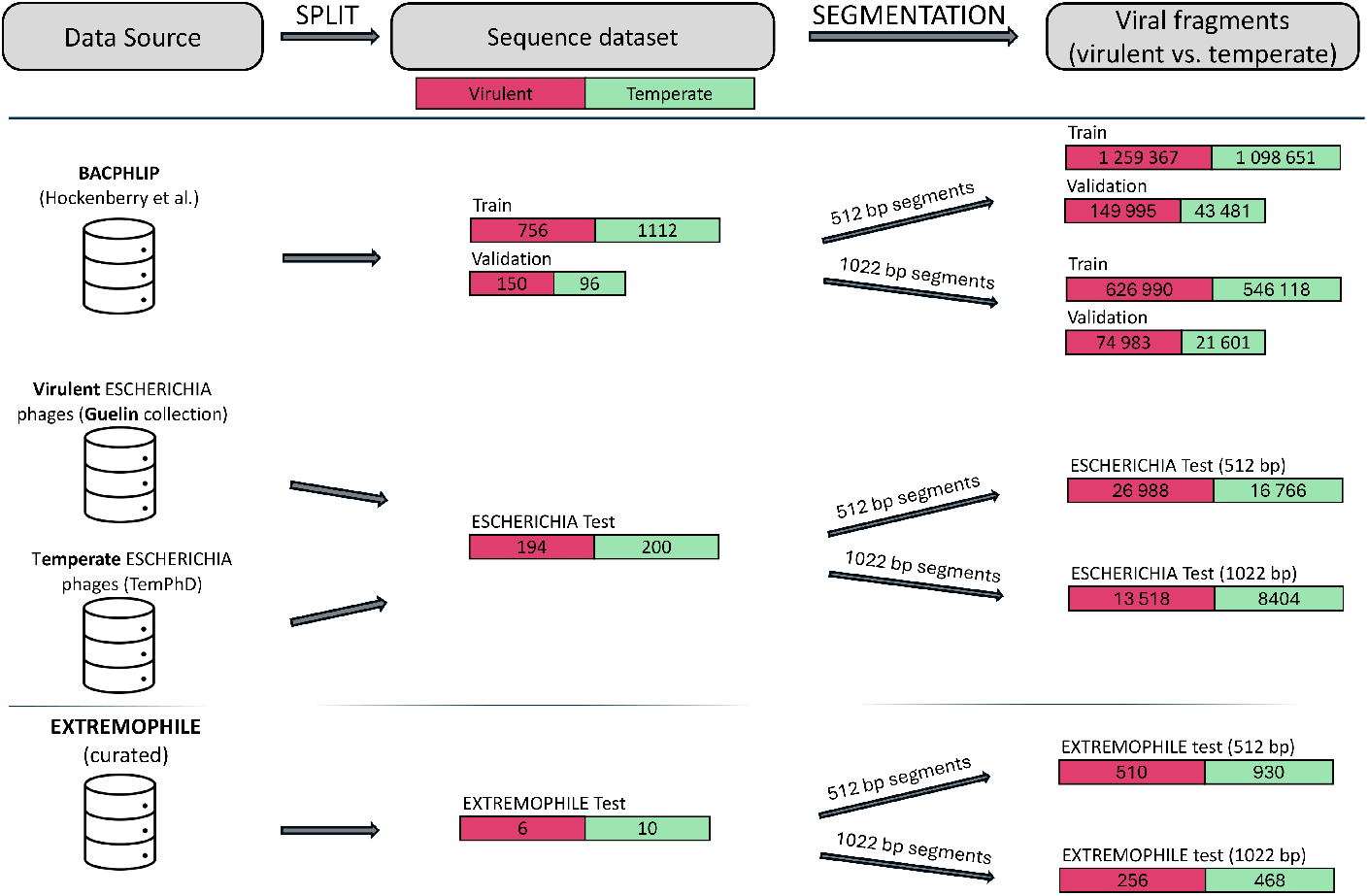
The datasets used in the study. Various independent data sources were used to train and evaluate the model performances (split). First, the annotated phage sequences are collected (sequence or contig datasets). Subsequently, the contigs are segmented into smaller 512 or 1022 bp length chunks. The numbers indicate the sizes of the datasets stratified by class. The training and validation datasets were collected from BACPHLIP dataset [20]. The *Escherichia* test consists of experimentally validated *Escherichia* phages (Guelin collection [35]) as well as high-quality, temperate phages downloaded from the TemPhD database ([41]). The Extremophile dataset was curated through a review of the literature.

To simulate these cases, we created two sets of short fragments, *L* = [512*bp*, 1022*bp*], by chunking the phage contigs into short reads, 512, and 1022bp length segments. We refer to this chunking process as segmentation and is described in detail in [33]. For the training and evaluation datasets, we applied random segmentations with expected coverages of 10x and 5x respectively, since in the case of fragmented assemblies not only the length is unknown, but we cannot be sure which part of the original contig is reported. On the other hand, a contiguous segmentation strategy was applied on the test set, because of its practicability.

In most practical cases of genome assemblies, the strand orientation is not known, thus we included reverse complement sequences for all genomic data.

### Ensuring dataset independence through similarity estimation

To ensure that the training and test sets do not overlap, we estimated the similarity between phage sequences using a MinHash algorithm. An all-against-all search was carried out to ensure that no phages are duplicated in the training and test sets, confirming their independence. The comparison was conducted at the nucleotide sequence level. Additionally, the MMSEQ2 [42] tool was applied on the sequence datasets. A lengthy signature containing 300 elements was used. These elements were derived from nucleotide sequences of length 21, using the MurmurHash3 hash function with a seed value of 3. Finally, the Jaccard coefficient was computed for each potential pair of sequences based on their respective signatures.

### Phage lifestyle prediction with genomic language models

Phage lifestyle prediction is formulated as a binary classification problem, similarly to the approaches of BACPHLIP, DeePhage, and PhaTYP. The task is to distinguish between virulent and non-virulent phage sequences, with temperate phages considered as the negative class.

The models used in this study, DNABERT-2, NT, and ProkBERT, are all encoder architectures. These models take tokenized sequences as input and produce vector representations as output. Classification is performed based on these output vectors. Training is conducted under a transfer learning paradigm; which is carried out as follows: a new model with a binary classification head is initialized from the pretrained model weights, and then the model weights are updated at each training iteration step to recognize virulent phage sequences. The models differ in various aspects such as size, tokenization strategy, and pretraining datasets.

### ProkBERT PhaStyle

The ProkBERT model family (`mini`, `mini-long`, and `mini-c`) accommodates different sequence vectorizations, tokenized with Local Context Aware (LCA) k-mers for efficient representation. Tokenization granularity varies across models, with `mini` and `mini-long` using 6-mers and `mini-c` relying on character-level tokens (see Figure 2 for an overview of the methodology).

**Fig. 2.**
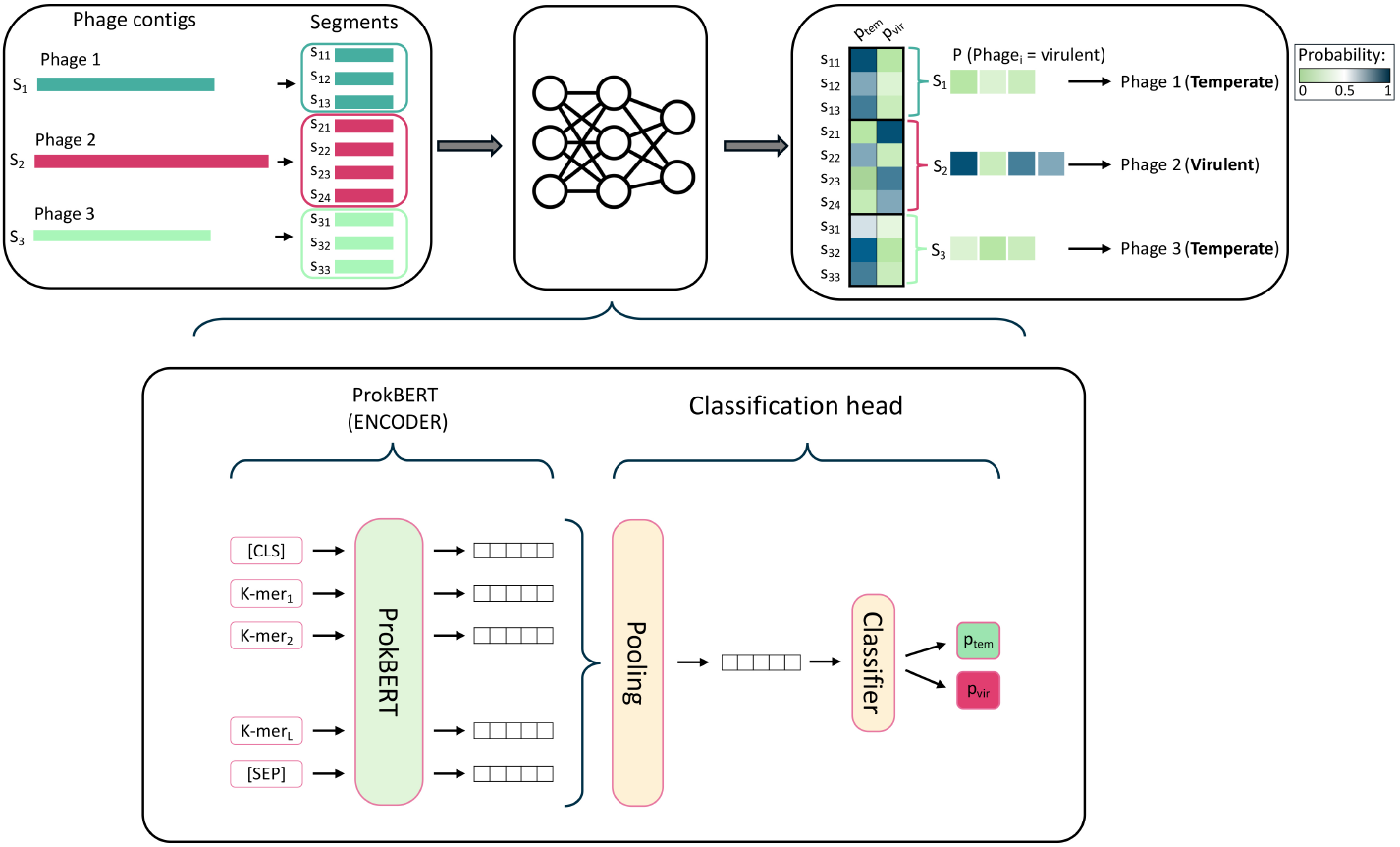
Methodology for Phage Lifestyle Prediction Using ProkBERT. The diagram illustrates the process of predicting phage lifestyles from nucleotide sequences. Phage contigs (S_1_, S_2_, S_3_) are segmented into smaller sequences (e.g., S_11_, S_12_, S_13_), which then are inputs to the ProkBERT encoder. The encoder processes these segments using k-mer tokenization and special tokens ([CLS], [SEP]) to generate hidden states. A classification head pools these representations to predict the probabilities of each segment being virulent (P_vir_) or temperate (P_tem_). Then the likelihood of each segment’s lifestyle is calculated. Finally, these probabilities are aggregated to classify the entire phage contig as either virulent or temperate.

Each model contains 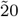M parameters, 6 layers, and 6 attention heads. The encoder generates token representations, which are pooled with learnable weights for classification (illustrated on Figure 2). For detailed architecture and classification steps, see supplementary materials.

The models were fine-tuned on the Komondor HPC system with AdamW optimization and adjusted batch sizes for 512 and 1022bp segments.

#### Weighted Voting for Contig Classification

To classify entire phage contigs, predictions from individual segments are combined using a weighted voting method, producing a single probability score per contig. See the supplementary material for the formalized approach.

#### Other genomic language models

DNABERT-2, a Transformer Encoder with 117M parameters, optimized for genome analysis, uses SentencePiece and BPE for DNA tokenization and incorporates ALiBi and Flash Attention for performance enhancement [43, 44, 45, 46]. Pretrained on genomes from 135 species (32.49 billion bases), it was fine-tuned for phage lifestyle prediction using the ‘AutoModelForSequenceClassification‘ class, batch sizes of 196 (512 bp) and 128 (1022 bp), and a learning rate of 0.00010.0001 with AdamW optimizer.

The Nucleotide Transformer was pretrained on the human reference genome and 850 additional genomes [30]. Available in 50M, 100M, and 500M parameter versions, it uses a 6-mer tokenization scheme with character-level adjustments. Fine-tuning used batch sizes adjusted to input and model sizes, and a learning rate of 0.00010.0001, with other parameters matching those for ProkBERT.

### Benchmarking

The performance of the new LM model-based approaches and current state-of-the-art methods was compared on phage lifestyle datasets: a) *Escherichia* phages and b) phages from extreme environments. The evaluation was conducted on both segment-level and full sequence datasets in order to simulate real-world applications where viral sequences are often fragmented. BACPHLIP, which only supports prediction on complete sequences, was evaluated accordingly. Additionally, we analyzed the inference speeds of the different approaches.

#### Out-of-sample prediction on Escherichia dataset

*Escherichia coli* is one of the best characterized bacteria and abundant experimental data is available about it, making it an ideal organism for validation purposes. We compared the methods on novel, experimentally validated virulent *Escherichia* phages. Although these phages are not in the training data, their close relatives are, making them good indicators of how well traditional models (PhaTYP, DeePhage, BACPHLIP) perform on representative cases. For genomic language models (ProkBERT, DNABERT-2, NT), the *Escherichia* sequences were treated as ’out-of-sample’ by discarding all phages infecting *Escherichia* from the training data, thus placing these models at a disadvantage compared to current methods.

First, we tested the models on phage fragments of various sizes, specifically 512 and 1022 bp. The results are in Table 1.

**Table 1.** Performance comparison of models on the *Escherichia* test set across various segment lengths. Best performances in each segment are highlighted in bold. The metrics in the table are as follows. Bal. Acc.: balanced accuracy, MCC: Matthews’ Correlation Coefficient, Spec.: specificity, Sens.: sensitivity

| Method | Bal. Acc. | MCC | Sens. | Spec. |
| --- | --- | --- | --- | --- |
| 512 bp |  |  |  |  |
| **ProkBERT-mini** | **0.91** | **0.83** | 0.94 | **0.89** |
| ProkBERT-mini-long | 0.90 | 0.82 | **0.96** | 0.85 |
| ProkBERT-mini-c | 0.89 | 0.80 | 0.95 | 0.84 |
| DNABERT-2-117M | 0.84 | 0.72 | 0.95 | 0.74 |
| Nuc.Trans.-50m | 0.85 | 0.72 | 0.92 | 0.78 |
| Nuc.Trans.-100m | 0.87 | 0.75 | 0.93 | 0.82 |
| Nuc.Trans.-500m | 0.88 | 0.78 | **0.96** | 0.80 |
| DeePhage | 0.86 | 0.71 | 0.84 | 0.88 |
| PhaTYP | **0.91** | **0.83** | 0.94 | 0.88 |
| 1022 bp |  |  |  |  |
| ProkBERT-mini | **0.94** | 0.88 | **0.97** | **0.91** |
| **ProkBERT-mini-long** | **0.94** | **0.89** | **0.97** | **0.91** |
| ProkBERT-mini-c | 0.93 | 0.87 | **0.97** | 0.89 |
| DNABERT-2-117M | 0.90 | 0.80 | 0.95 | 0.85 |
| Nuc.Trans.-50m | 0.90 | 0.80 | 0.94 | 0.85 |
| Nuc.Trans.-100m | 0.92 | 0.83 | 0.94 | 0.89 |
| Nuc.Trans.-500m | 0.91 | 0.84 | 0.96 | 0.87 |
| DeePhage | 0.91 | 0.82 | 0.94 | 0.88 |
| PhaTYP | 0.92 | 0.84 | 0.96 | 0.87 |

Generally speaking, both Genomic LMs and current computational approaches performed well on longer segments, with somewhat larger spread on shorter fragments which are more difficult to classify. Prokbert models came out consistently at the top of the list in both cases, with a 0.91 and 0.94 balanced accuracy being the highest result, respectively. It is conspicuous that all models performed better on 1022 base pairs. `ProkBERT-mini` and `ProkBERT-mini-long` both achieved the highest balanced accuracy (0.94). A more detailed summary of the results can be found in Supplementary Table 6.

A comparison of *Escherichia* full sequences, including BACPHLIP (Supplementary Table 3), was performed. Genomic LMs’ predictions for contigs were based on the weighted average of 512 bp segment predictions (see Methods). In this comparison, ProkBERT and all genomic LMs consistently performed well, each reaching top metrics with an MCC of 0.98. BACPHLIP, DeePhage, and PhaTYP achieved MCC values of 0.96, 0.95, and 0.88, respectively.

These results suggest that current computational approaches are reliable for phages well-represented in their training data. The ProkBERT models, particularly, showed good generalization capabilities, maintaining high performance even when trained on different data distributions.

#### Performance on Extremophile dataset

Phages found in extreme environments such as the deep sea, high pressure zones, cold climates, and acidic conditions, are less well characterized than those of the human gut. Such environmental phages are generally challenging and often underrepresented in databases, therefore they serve as good indicators of the model performances. The results of this analysis are summarized in Table 2.

**Table 2.** Performance comparison of models on the Extremophile test set across various segment lengths. Best performances in each segment are highlighted in bold. The metrics in the table are as follows. Bal. Acc.: balanced accuracy, MCC: Matthews’ Correlation Coefficient, Spec.: specificity, Sens.: sensitivity

| Method | Bal. Acc. | MCC | Sens. | Spec. |
| --- | --- | --- | --- | --- |
| 512 bp |  |  |  |  |
| **ProkBERT-mini** | **0.93** | **0.83** | 0.99 | 0.87 |
| **ProkBERT-mini-long** | **0.93** | 0.82 | **1** | 0.86 |
| ProkBERT-mini-c | 0.92 | 0.80 | 0.99 | 0.84 |
| DNABERT-2-117M | 0.89 | 0.74 | 0.99 | 0.79 |
| Nuc.Trans.-50m | 0.91 | 0.79 | 0.98 | 0.84 |
| Nuc.Trans.-100m | 0.90 | 0.76 | 0.97 | 0.82 |
| Nuc.Trans.-500m | 0.91 | 0.78 | 0.99 | 0.82 |
| DeePhage | 0.87 | 0.75 | 0.84 | **0.91** |
| PhaTYP | 0.76 | 0.52 | 0.74 | 0.79 |
| 1022 bp |  |  |  |  |
| **ProkBERT-mini** | **0.96** | **0.91** | **1** | **0.93** |
| **ProkBERT-mini-long** | **0.96** | 0.90 | **1** | 0.92 |
| ProkBERT-mini-c | 0.94 | 0.86 | **1** | 0.89 |
| DNABERT-2-117M | 0.94 | 0.85 | 0.98 | 0.90 |
| Nuc.Trans.-50m | 0.93 | 0.83 | 0.99 | 0.87 |
| Nuc.Trans.-100m | 0.95 | 0.88 | 0.98 | 0.91 |
| Nuc.Trans.-500m | 0.96 | 0.89 | 1 | 0.91 |
| DeePhage | 0.92 | 0.80 | 0.96 | 0.87 |
| PhaTYP | 0.80 | 0.58 | 0.84 | 0.76 |

Genomic LMs performed better on this task than current SOTA methods. Similar tendencies were observed as in the previous task (*Escherichia* dataset). All methods performed better when larger contextual information was available. ProkBERT models consistently ranked at the top in both fragment sizes. A more detailed summary of the results with more metrics can be found in Supplementary Table 7.

A comparison on Extremophile full sequences (Supplementary Table 4) revealed that all GNLMs and also DeePhage showed perfect performance (*MCC* = 1.00); BACPHLIP and PhaTYP produced MCC values of 0.77 and 0.52, respectively.

In summary, the two datasets (*Escherichia* and Extremophile) were selected so as to test the models’ performance and generalization capability. Interestingly, despite being placed at a disadvantage compared to PhaTYP, DeePhage and BACPHLIP that were trained on a broader dataset, the genomic LMs showed unexpectedly good prediction performance on both datasets. One reason for this may be that, during the pretraining phase, they capture patterns that are useful for this task. Apparently, ProkBERT models consistently outperformed other models across various datasets and segment lengths. This is particularly notable since DNABERT-2 and NT models have much larger capacities (2-20 times more parameters) compared to ProkBERT. We believe this is due to the fact that ProkBERT was pretrained on a much larger microbial dataset and employs LCA tokenization, which provides more robust vectorization of nucleotide sequences.

In conclusion, ProkBERT PhaStyle is the fine-tuned version of ProkBERT-mini. The `mini` model was selected due to its consistent and robust performance across various scenarios.

#### Inference speed and running times

The computational performance of the models was assessed using a set of 1,000 randomly selected sequences from the BACPHLIP dataset. The evaluations were conducted on identical hardware configurations (Supplementary Figure 1). Among the genomic language models, `ProkBERT-mini-long` was the fastest with execution time of 132 sec and the highest inference speed of 0.52 MB/sec. `mini` and `mini-c` performed similarly, with slightly longer execution times of 141 and 146 seconds, and inference speeds of 0.49 and 0.47 MB/sec, respectively.

As expected, larger transformer models such as the DNABERT-2 (117m parameters) and NT (500m parameters) require 2x-3.8x more time for to evaluate samples. Since these models apply different tokenization, the input size (number of tokens used in the representations) signigicantly influences the performance, i.e. the less token the faster the prediction is.

DeePhage and PhaTYP displayed longer execution times and lower speeds compared to the genomic language models. DeePhage executed in 159 seconds with a prediction rate of 0.43 MB/sec. The database search based approaches (PhaTYP and BACPHLIP) required significantly more time (2718 and 7125 second, respectively).^1^

### Limitations and practical implications

Our proposed method, like existing phage lifestyle prediction tools, assumes input sequences are known to be viral. Both the genomic language models and SOTA methods compared in this work are binary classifiers that require pre-identified phage sequences as input, labeling them as either virulent or temperate. Consequently, an upstream viral sequence identification step is necessary to distinguish phage sequences from non-phage sequences in metagenomic datasets.

To address this requirement, several viral identification tools can be integrated into the analysis pipeline, including VirSorter2 [47], VIBRANT [48], DeepVirFinder [49], and ProkBERT [33], which is particularly effective on short fragments. For thorough reviews of these tools, see [50, 51, 52]. Another consideration is the computational resources required for model inference. Most of the models presented in this work, with the exception of BACPHLIP, perform best with GPU support due to the computational demands of deep learning models. While this may pose a challenge for some users, GPU resources are increasingly accessible. Cloud-based platforms like Google Colab offer free GPU access, and many institutions provide computational resources to researchers.

## Conclusion

This study introduces and evaluates a novel approach for phage lifestyle prediction by applying pretrained genomic language models and fine-tuning them for this specific task. The ability of these models, particularly ProkBERT, to directly process nucleotide sequences without the need for protein annotation or complex database searches is significant. This approach reduces biases and computational overhead.

ProkBERT, designed specifically for microbial classification tasks, consistently outperformed current computational approaches and other universal language models such as Nucleotide Transformer and DNABERT-2. The ProkBERT PhaStyle models proved to be simple, easy to train, and fast, demonstrating high efficiency and accuracy, even when classifying short sequence fragments. This underscores their applicability to metagenomic and metavirome datasets, where sequences are often fragmented.

Furthermore, the new models exhibited good generalization capabilities, performing well on previously unseen phage sequences, including those from extreme environments. This robustness and versatility make ProkBERT a valuable tool for phage lifestyle prediction in diverse ecological and clinical settings. Future research should aim to increase the interpretability of these models and explore their broader applications in microbial genomics.

## Key points

We demonstrated that complex machine learning task such as phage lifestyle prediction can be solved directly from nucleotide sequences without the need for complex bioinformatics pipelines or manually curated protein annotations.

We showed that genomic language models and the transfer learning paradigm offer a simple, but efficient alternative to database search based methods. They offer better generalization, while being faster and more accurate.

We introduced ProkBERT PhaStyle, which is compact and consinstently outperformed alternative genomic language models and alternative methods in various challanging scenarios, such as classifing short fragments, extremophile phage sequences and unseen phage sequences.

## Data availability

The datasets generated and analyzed for this study are available on Zenodo: 10.5281/zenodo.13959905 The various containerized versions of BACPHLIP, DNABERT-2, DeePhage,Nucleotide Transformers and ProkBERT PhaStyle are available on Zenodo (Apptainer) and on Docker Hub https://hub.docker.com/repository/docker/obalasz/phage/general

## Code availability

The code associated with this study is available on GitHub: https://github.com/nbrg-ppcu/PhaStyle Additionally, datasets, models and codes can be accessed at the following repositories: GitHub – https://github.com/nbrg-ppcu/PhaStyle, HuggingFace –https://huggingface.co/neuralbioinfo.

## Author contributions statement

JJ (Judit Juhász): Data analysis, Assessments, Data curation, Methodology, Validation, Writing. BB: Data curation, Investigation, Methodology, Software JJ (János Juhász): Data curation, Interpretation, Writing. LNN: Formal analysis, Visualization, Writing, Analysis. SP: Conceptualization, Review writing. BL: Conceptualization, Data curation, Formal analysis, Funding acquisition, Investigation, Methodology, Project administration, Resources, Software, Supervision, Validation, Visualization, Writing.

## Acknowledgments

The authors gratefully acknowledge the HPC RIVR consortium (www.hpc-rivr.si) and EuroHPC JU (`eurohpc-ju.europa.eu`) for funding this research by providing computing resources of the HPC system Vega at the Institute of Information Science (www.izum.si) as well as to HPC-KIFU Komondor and LUMI.

## Footnotes

1 Supplementary Table 5 contains the data on execution time and inference speed.

## References

1. Cristina Howard-Varona, Katherine R Hargreaves, Stephen T Abedon, and Matthew B Sullivan. Lysogeny in nature: mechanisms, impact and ecology of temperate phages. The ISME journal, 11(7):1511–1520, 2017.

2. Zhirui Cao, Naoki Sugimura, Elke Burgermeister, Matthias P Ebert, Tao Zuo, and Ping Lan. The gut virome: A new microbiome component in health and disease. EBioMedicine, 81, 2022.

3. Mohammadali Khan Mirzaei and Li Deng. New technologies for developing phage-based tools to manipulate the human microbiome. Trends in microbiology, 30(2):131–142, 2022.

4. Steffanie A Strathdee, Graham F Hatfull, Vivek K Mutalik, and Robert T Schooley. Phage therapy: From biological mechanisms to future directions. Cell, 186(1):17–31, 2023.

5. Beatriz Gutiérrez and Pilar Domingo-Calap. Phage therapy in gastrointestinal diseases. Microorganisms, 8(9):1420, 2020.

6. Yashoda Bhattarai and Purna C. Kashyap. Engineering human microbiota: influencing cellular and community stability. Nature Reviews Gastroenterology & Hepatology, 17:297–309, 2020.

7. Jee Loon Foo, Hua Ling, Yung Seng Lee, and Matthew Wook Chang. Microbiome engineering: Current applications and its future. Biotechnology journal, 12(3):1600099, 2017.

8. Jorge A Moura de Sousa, Eugen Pfeifer, Marie Touchon, and Eduardo PC Rocha. Causes and consequences of bacteriophage diversification via genetic exchanges across lifestyles and bacterial taxa. Molecular biology and evolution, 38(6):2497–2512, 2021.

9. Andrey N Shkoporov, Adam G Clooney, Thomas DS Sutton, Feargal J Ryan, Karen M Daly, James A Nolan, Siobhan A McDonnell, Ekaterina V Khokhlova, Lorraine A Draper, Amanda Forde, et al. The human gut virome is highly diverse, stable, and individual specific. Cell host & microbe, 26(4):527–541, 2019.

10. Katelyn McNair, Barbara A Bailey, and Robert A Edwards. Phacts, a computational approach to classifying the lifestyle of phages. Bioinformatics, 28(5):614–618, 2012.

11. Adam J. Hockenberry and Claus O. Wilke. BACPHLIP: Predicting Bacteriophage Lifestyle from Conserved Protein Domains. PeerJ, 9:e11396, 2021.

12. Jiayu Shang, Xubo Tang, and Yanni Sun. Phatyp: predicting the lifestyle for bacteriophages using bert. Briefings in Bioinformatics, 24(1):bbac487, 2023.

13. Kai Song. Classifying the lifestyle of metagenomically-derived phages sequences using alignment-free methods. Frontiers in microbiology, 11:567769, 2020.

14. Shufang Wu, Zhencheng Fang, Jie Tan, Mo Li, Chunhui Wang, Qian Guo, Congmin Xu, Xiaoqing Jiang, and Huaiqiu Zhu. Deephage: distinguishing virulent and temperate phage-derived sequences in metavirome data with a deep learning approach. Gigascience, 10(9):giab056, 2021.

15. David Paez-Espino, Emiley A Eloe-Fadrosh, Georgios A Pavlopoulos, Alex D Thomas, Marcel Huntemann, Natalia Mikhailova, Edward Rubin, Natalia N Ivanova, and Nikos C Kyrpides. Uncovering Earth’s virome. Nature, 536(7617):425–430, 2016.

16. Luis F Camarillo-Guerrero, Alexandre Almeida, Guillermo Rangel-Pineros, Robert D Finn, and Trevor D Lawley. Massive expansion of human gut bacteriophage diversity. Cell, 184(4):1098–1109, 2021.

17. Antonio Pedro Camargo, Stephen Nayfach, I-Min A Chen, Krishnaveni Palaniappan, Anna Ratner, Ken Chu, Stephan J Ritter, TBK Reddy, Supratim Mukherjee, Frederik Schulz, et al. IMG/VR v4: an expanded database of uncultivated virus genomes within a framework of extensive functional, taxonomic, and ecological metadata. Nucleic acids research, 51(D1):D733–D743, 2023.

18. Stephen Nayfach, David Páez-Espino, Lee Call, Soo Jen Low, Hila Sberro, Natalia N Ivanova, Amy D Proal, Michael A Fischbach, Ami S Bhatt, Philip Hugenholtz, et al. Metagenomic compendium of 189,680 DNA viruses from the human gut microbiome. Nature microbiology, 6(7):960–970, 2021.

19. Shiraz A Shah, Ling Deng, Jonathan Thorsen, Anders G Pedersen, Moïra B Dion, Josué L Castro-Mejía, Ronalds Silins, Fie O Romme, Romain Sausset, Leon E Jessen, et al. Expanding known viral diversity in the healthy infant gut. Nature microbiology, 8(5):986–998, 2023.

20. Adam J Hockenberry and Claus O Wilke. BACPHLIP: predicting bacteriophage lifestyle from conserved protein domains. PeerJ, 9:e11396, 2021.

21. Ashish Vaswani, Noam Shazeer, Niki Parmar, Jakob Uszkoreit, Llion Jones, Aidan N Gomez, Lukasz Kaiser, and Illia Polosukhin. Attention is all you need. Advances in neural information processing systems, 30, 2017.

22. Roshan Rao, Jason Liu, Robert Verkuil, Joshua Meier, John F. Canny, Pieter Abbeel, Tom Sercu, and Alexander Rives. MSA Transformer. bioRxiv, 2021.

23. Zeming Lin, Halil Akin, Roshan Rao, Brian Hie, Zhongkai Zhu, Wenting Lu, Nikita Smetanin, Allan dos Santos Costa, Maryam Fazel-Zarandi, Tom Sercu, Sal Candido, et al. Language models of protein sequences at the scale of evolution enable accurate structure prediction. bioRxiv, 2022.

24. Alexander Rives, Joshua Meier, Tom Sercu, Siddharth Goyal, Zeming Lin, Jason Liu, Demi Guo, Myle Ott, C Lawrence Zitnick, Jerry Ma, et al. Biological structure and function emerge from scaling unsupervised learning to 250 million protein sequences. Proceedings of the National Academy of Sciences, 118(15):e2016239118, 2021.

25. John Jumper, Richard Evans, Alexander Pritzel, Tim Green, Michael Figurnov, Olaf Ronneberger, Kathryn Tunyasuvunakool, Russ Bates, Augustin Žídek, Anna Potapenko, et al. Highly accurate protein structure prediction with AlphaFold. Nature, 596(7873):583–589, 2021.

26. Gustaf Ahdritz, Nazim Bouatta, Christina Floristean, Sachin Kadyan, Qinghui Xia, William Gerecke, Timothy J O’Donnell, Daniel Berenberg, Ian Fisk, Niccoló Zanichelli et al. OpenFold: Retraining AlphaFold2 yields new insights into its learning mechanisms and capacity for generalization. Nature Methods, pages 1–11, 2024.

27. Yanrong Ji, Zhihan Zhou, Han Liu, and Ramana V Davuluri. DNABERT: pre-trained Bidirectional Encoder Representations from Transformers model for DNA-language in genome. Bioinformatics, 37(15):2112–2120, 2021.

28. Zhihan Zhou, Yanrong Ji, Weijian Li, Pratik Dutta, Ramana Davuluri, and Han Liu. Dnabert-2: Efficient foundation model and benchmark for multi-species genome. arXiv preprint 2306.15006, 2023.

29. Veniamin Fishman, Yuri Kuratov, Maxim Petrov, Aleksei Shmelev, Denis Shepelin, Nikolay Chekanov, Olga Kardymon, and Mikhail Burtsev. GENA-LM: A Family of Open-Source Foundational Models for Long DNA Sequences. bioRxiv. 2023.

30. Marie Lopez, Hugo Dalla-Torre, Liam Gonzalez, Javier Mendoza-Revilla, Nicolas Lopez Carranza, Adam Grzywaczewski, Francesco Oteri, Christian Dallago, Evan Trop, Hassan Sirelkhatim, et al. The Nucleotide Transformer: Building and Evaluating Robust Foundation Models for Human Genomics. 2023.

31. Albert Gu and Tri Dao. Mamba: Linear-time sequence modeling with selective state spaces. arXiv preprint 2312.00752, 2023.

32. Eric Nguyen, Michael Poli, Marjan Faizi, Armin Thomas, Michael Wornow, Callum Birch-Sykes, Stefano Massaroli, Aman Patel, Clayton Rabideau, Yoshua Bengio, et al. HyenaDNA: Long-range genomic sequence modeling at single nucleotide resolution. Advances in neural information processing systems, 36, 2024.

33. Balázs Ligeti, István Szepesi-Nagy, Babett Bodnár, Noémi Ligeti-Nagy, and János Juhász. ProkBERT family: genomic language models for microbiome applications. Frontiers in Microbiology, 14:1331233, 2024.

34. Travis N. Mavrich and Graham F. Hatfull. A large dataset of phage genomes for bacteriophage lifestyle prediction. mBio, 8(3):e02133–17, 2017.

35. J.P. Gaborieau et al. A Collection of Taxonomically Diverse Escherichia Bacteriophages: Insights into Virulent and Temperate Interactions. bioRxiv, 2023.

36. Katarzyna Bujak, Przemyslaw Decewicz, Michal Kitowicz, and Monika Radlinska. Characterization of three novel virulent Aeromonas phages provides insights into the diversity of the Autographiviridae family. Viruses, 14(5):1016, 2022.

37. Tim Engelhardt, Monika Sahlberg, Heribert Cypionka, and Bert Engelen. Biogeography of Rhizobium radiobacter and distribution of associated temperate phages in deep subseafloor sediments. The ISME journal, 7(1):199–209, 2013.

38. Mitsuhiro Yoshida, Yukari Yoshida-Takashima, Takuro Nunoura, and Ken Takai. Genomic characterization of a temperate phage of the psychrotolerant deep-sea bacterium Aurantimonas sp. Extremophiles, 19:49–58, 2015.

39. Yue Su, Wenjing Zhang, Yantao Liang, Hongmin Wang, Yundan Liu, Kaiyang Zheng, Ziqi Liu, Hao Yu, Linyi Ren, Hongbing Shao, et al. Identification and genomic analysis of temperate Halomonas bacteriophage vB HmeY H4907 from the surface sediment of the Mariana Trench at a depth of 8,900 m. Microbiology Spectrum, 11(5):e01912–23, 2023.

40. Pablo Tapia, Francisco Moya Flores, Paulo C Covarrubias, Lillian G Acunã, David S Holmes, and Raquel Quatrini. Complete Genome Sequence of Temperate Bacteriophage Aca ML1 from the Extreme Acidophile Acidithiobacillus caldus ATCC 51756. 2012.

41. Xianglilan Zhang, Ruohan Wang, Xiangcheng Xie, Yunjia Hu, Jianping Wang, Qiang Sun, Xikang Feng, Wei Lin, Shanwei Tong, Wei Yan, et al. Mining bacterial NGS data vastly expands the complete genomes of temperate phages. NAR Genomics and Bioinformatics, 4(3):lqac057, 2022.

42. Martin Steinegger and Johannes Söding. Mmseqs2 enables sensitive protein sequence searching for the analysis of massive data sets. Nature biotechnology, 35(11):1026–1028, 2017.

43. Taku Kudo and John Richardson. Sentencepiece: A simple and language independent subword tokenizer and detokenizer for neural text processing. In Proceedings of the 2018 Conference on Empirical Methods in Natural Language Processing: System Demonstrations, pages 66– 71, 2018.

44. Rico Sennrich, Barry Haddow, and Alexandra Birch. Neural machine translation of rare words with subword units. arXiv preprint 1609.08144, 2016.

45. Ofir Press, Noah A Smith, and Mike Lewis. Train short, test long: Attention with linear biases enables input length extrapolation. arXiv preprint 2108.12409, 2021.

46. Tri Dao, Daniel Y Fu, Stefano Ermon, Atri Rudra, and Christopher Ré. Flashattention: Fast and memory-efficient exact attention with io-awareness. In Proceedings of the IEEE/CVF Conference on Computer Vision and Pattern Recognition, pages 12785–12795, 2022.

47. Jiarong Guo, Ben Bolduc, Ahmed A Zayed, Arvind Varsani, Guillermo Dominguez-Huerta, Tom O Delmont, Akbar Adjie Pratama, M Consuelo Gazituá, Dean Vik, Matthew B Sullivan, et al. VirSorter2: a multi-classifier, expert-guided approach to detect diverse DNA and RNA viruses. Microbiome, 9:1–13, 2021.

48. Kristopher Kieft, Zhichao Zhou, and Karthik Anantharaman. VIBRANT: automated recovery, annotation and curation of microbial viruses, and evaluation of viral community function from genomic sequences. Microbiome, 8:1–23, 2020.

49. Jie Ren, Kai Song, Chao Deng, Nathan A Ahlgren, Jed A Fuhrman, Yi Li, Xiaohui Xie, Ryan Poplin, and Fengzhu Sun. Identifying viruses from metagenomic data using deep learning. Quantitative Biology, 8(1):64–77, 2020.

50. Ling-Yi Wu, Yasas Wijesekara, Gonçalo J Piedade, Nikolaos Pappas, Corina PD Brussaard, and Bas E Dutilh. Benchmarking bioinformatic virus identification tools using real-world metagenomic data across biomes. Genome Biology, 25(1):97, 2024.

51. Kenneth E Schackart III, Jessica B Graham, Alise J Ponsero, and Bonnie L Hurwitz. Evaluation of computational phage detection tools for metagenomic datasets. Frontiers in Microbiology, 14:1078760, 2023.

52. Siu Fung Stanley Ho, Nicole E Wheeler, Andrew D Millard, and Willem van Schaik. Gauge your phage: benchmarking of bacteriophage identification tools in metagenomic sequencing data. Microbiome, 11(1):84, 2023.

