## Supplementary material for "ProkBERT PhaStyle: Accurate Phage Lifestyle Prediction with Pretrained Genomic Language Models"

Judit Juhász et al.

#### ProkBERT PhaStyle model overview and methodology

The ProkBERT model family, comprising `mini`, `mini-long`, and `mini-c`, is designed to accommodate different modes of sequence vectorization with varying maximum input lengths. Tokenization is performed using Local Context Aware (LCA) k-mers. For instance, LCA tokenization of a sequence like ATACGGT with k-mer= 3 and shift= 2 results in tokens $\{ATA, ACG, GGT\}$. This approach allows for the representation of longer sequences with fewer tokens as the shift value increases, although it comes at the cost of increasing the number of potential vectors. Specifically, the `mini-c` model uses character-level tokenization (k-mer= 1, shift= 1), while the `mini` and `mini-long` models utilize overlapping 6-mers with shifts of 1 and 2, respectively.

Each model in the ProkBERT family (`mini`, `mini-long`, and `mini-c`) contains approximately 20 million parameters, with 6 layers, 6 attention heads, and an embedding size of 386.

For classification, we used a simple model previously introduced in [1]. Briefly, the encoder (ProkBERT) generates representations (last hidden states) for each token, which are pooled with learnable weights, and classification is performed based on the pooled vector.

More formally, given an input sequence $\mathbf{X} = \{x_1, x_2, ..., x_n\}$, where $n$ is the sequence length, the ProkBERT base model first transforms $\mathbf{X}$ into a sequence of hidden states $\mathbf{H} = \{h_1, h_2, ..., h_n\}$, each $h_i \in \mathbb{R}^d$, with $d$ representing the hidden size. The hidden states $\mathbf{H}$ are then processed through a weighting layer, which assigns a weight $w_i$ to each token's hidden state $h_i$:

$$w_i = \text{softmax}\left(\mathbf{W}_w h_i + b_w\right)$$

where $\mathbf{W}_w \in \mathbb{R}^{1\times d}$ and $b_w \in \mathbb{R}$ are the parameters of the weighting layer, and the softmax function is applied across the sequence length dimension to ensure that the weights sum to 1.

Following the assignment of weights, a weighted sum pooling operation is performed to aggregate the sequence into a single vector $\mathbf{p} \in \mathbb{R}^d$:

$$\mathbf{p} = \sum_{i=1}^{n} w_i h_i$$

The pooled output $\mathbf{p}$ is then passed through a dropout layer for regularization, followed by a linear classifier to produce the final logits $\mathbf{z} \in \mathbb{R}^2$ for the two classes:

$$\mathbf{z} = \mathbf{W}_c(\text{dropout}(\mathbf{p})) + b_c$$

where $\mathbf{W}_c \in \mathbb{R}^{2\times d}$ and $b_c \in \mathbb{R}^2$ are the parameters of the classifier.

The model outputs logits $\mathbf{z}$, from which the class probabilities can be derived using the softmax function:

$$\mathbf{y} = \text{softmax}(\mathbf{z})$$

Optimization during training involved adjusting batch sizes and gradient accumulation steps based on the model configuration and sequence length. For `mini` and `mini-c` the batch sizes (per device) were 336 and 192 for 512 and 1022bp long segments, respectively. The batch sizes for `mini-long` were chosen as 512 and 256 for the two segment lengths. The AdamW optimizer was used with learning rates of 0.0001 and 0.0005, $\beta_1 = 0.95$, $\beta_2 = 0.98$, weight decay of 0.1, and $\epsilon = 5 \times 10^{-5}$. Pytorch's cross entropy loss was used. The models were fine-tuned on the Komondor High Performance Computing (HPC) system, equipped with AMD EPYC 7763 64-Core Processors and 4 NVIDIA A100 (40GB) Tensor Core GPUs per node. The software stack included PyTorch version 2.0.1+cu117 and the Huggingface Transformers library version 4.33.2.

**Weighted voting procedure for contigs** The process of weighted voting in the context of predicting the lifestyle of bacteriophages, based on predictions made on individual segments, is designed to aggregate segment-level predictions into a single prediction for the entire phage contig. Given a phage contig divided into multiple segments, the model predicts the probability of each segment being virulent. These probabilities are then aggregated to make a final prediction for the whole contig. The process is mathematically formalized as follows:

Given a dataset $\mathcal{D}$ consisting of tuples $(s, p_0, p_1)$ for each segment, where $s$ is the segment identifier, $p_0$ is the probability of the segment belonging to class 0 (non-virulent), and $p_1$ is the probability of the segment belonging to class 1 (virulent), the goal is to compute a consolidated prediction for each unique sequence (phage contig) represented by multiple segments.

The procedure is as follows:

1. **Mean Probability Calculation:** For each unique sequence identifier $i$, calculate the mean probabilities for each class across all segments belonging to the same sequence. This is given by:

$$\bar{p}_0^{(i)} = \frac{1}{N_i} \sum_{j=1}^{N_i} p_0^{(j)}, \quad \bar{p}_1^{(i)} = \frac{1}{N_i} \sum_{j=1}^{N_i} p_1^{(j)}$$

where $N_i$ is the number of segments for sequence $i$, and $p_0^{(j)}$, $p_1^{(j)}$ are the probabilities for the $j^{th}$ segment of being non-virulent and virulent, respectively.

2. **Class Prediction:** The predicted class for each sequence $i$ is determined by the class with the higher mean probability:

$$y_{\text{pred}}^{(i)} = \begin{cases} 0 & \text{if } \bar{p}_0^{(i)} > \bar{p}_1^{(i)} \\ 1 & \text{otherwise} \end{cases}$$

This weighted voting approach allows for the integration of predictions from multiple segments.

### Other genomic language models

#### DNABERT-2

DNABERT-2 is a Transformer Encoder model with 117 million parameters, optimized for genomic sequence analysis. The model uses SentencePiece [2] combined with Byte Pair Encoding (BPE) [3] for tokenizing DNA sequences, enabling flexible handling of nucleotide sequences. DNABERT-2 was pretrained on a diverse genomic dataset, including genomes from 135 species, covering a total of 32.49 billion nucleotide bases. The model incorporates Attention with Linear Biases (ALiBi) [4] and Flash Attention [5] to enhance performance, particularly in terms of memory efficiency and speed for long-sequence processing.

For consistent deployment, DNABERT-2 was packaged in an Apptainer container configured with Python 3.8, PyTorch 1.13, and Transformers 4.29.2, ensuring compatibility with the original model settings.

Fine-tuning DNABERT-2 for phage lifestyle prediction involved the ‘AutoModelForSequenceClassification‘ class. Batch sizes of 196 for 512-base sequences and 128 for 1022-base sequences were used, with training run for two epochs. The AdamW optimizer was applied with hyperparameters $\beta_1 = 0.95$, $\beta_2 = 0.98$, weight decay of 0.1, $\epsilon = 5 \times 10^{-5}$, and a learning rate of 0.0001, chosen based on optimal results reported by the original authors.

##### Nucleotide Transformer

The Nucleotide Transformer was pretrained on two distinct datasets: the human reference genome, which includes 3,202 diverse human genomes, and a set of 850 genomes from various species [6]. This model is available in multiple configurations: nucleotide-transformer-v2-50m-multi-species, nucleotide-transformer-v2-100m-multi-species, and nucleotide-transformer-v2-500m-multi-species. Tokenization was applied using a non-overlapping 6-mer scheme, with a fallback to character-level tokenization for frames containing non-standard bases (e.g., characters other than A, C, T, G).

Fine-tuning the Nucleotide Transformer used batch sizes adjusted for input and model size, and two learning rates were tested: 0.0001 and 0.00005, with the former providing superior performance. Fine-tuning for binary classification tasks was achieved using the ‘AutoModelForSequenceClassification‘ class. Other training parameters, including the software and hardware environments, mirrored those used for the ProkBERT fine-tuning, ensuring consistency across models.

#### Evaluation metrics and definitions

**MCC (Matthews Correlation Coefficient)**: Used for binary classifications, defined as:

$$\text{MCC} = \frac{TP \times TN - FP \times FN}{\sqrt{(TP + FP)(TP + FN)(TN + FP)(TN + FN)}}$$

**F1 Score**: The harmonic mean of precision and recall, given by:

$$F1 = 2 \times \frac{\text{Precision} \times \text{Recall}}{\text{Precision} + \text{Recall}}$$

**Accuracy**: Represents the proportion of correctly predicted instances to the total, defined as:

$$\text{Accuracy} = \frac{TP + TN}{TP + TN + FP + FN}$$

**Balanced Accuracy**: An average of recall obtained on each class, giving equal weight to the performance on each class, defined as:

$$\text{Balanced Accuracy} = \frac{\text{Sensitivity} + \text{Specificity}}{2}$$

**Sensitivity (Recall)**: The proportion of actual positives correctly identified:

$$\text{Sensitivity} = \frac{TP}{TP + FN}$$

**Specificity**: The proportion of actual negatives correctly identified:

$$\text{Specificity} = \frac{TN}{TN + FP}$$

**Cross Entropy Loss**: A measure of the difference between two probability distributions for a given random variable or set of events. In binary classification, it is defined as:

$$\text{Cross Entropy Loss} = -\frac{1}{N}\sum_{i=1}^{N}[y_i \log(\hat{y}_i) + (1 - y_i)\log(1 - \hat{y}_i)]$$

where $N$ is the number of observations, $y_i$ is the actual value, and $\hat{y}_i$ is the predicted probability of the instance being in class 1.

#### Supplementary Figures

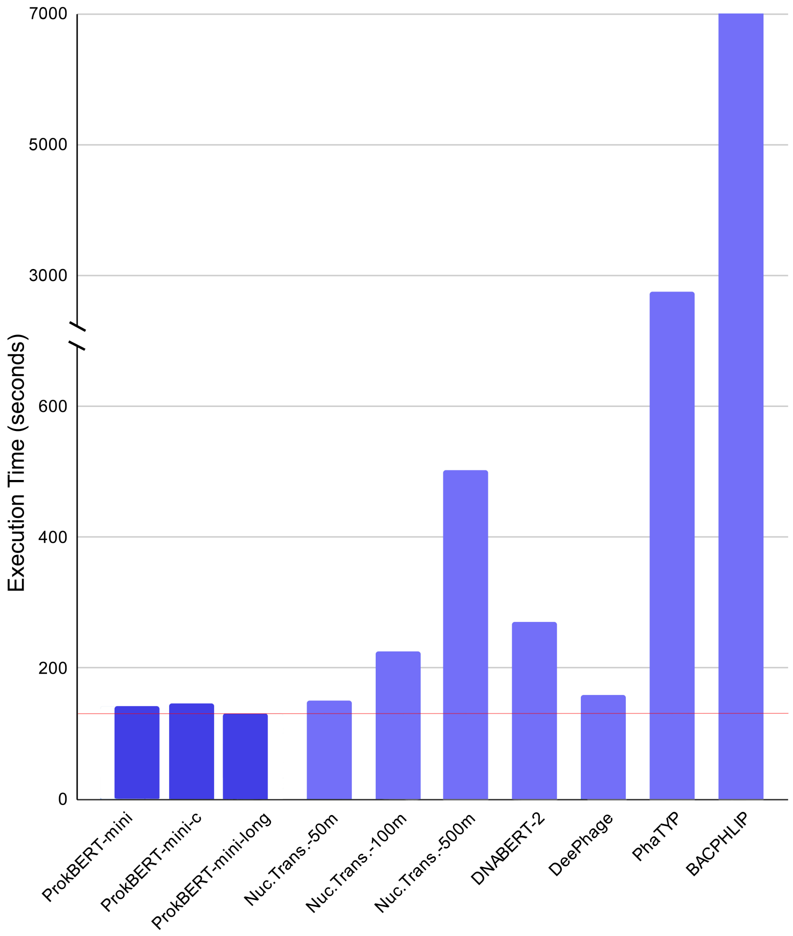


Figure 1: Execution times of various models for phage lifestyle prediction. The $y$ axis represents the execution time in seconds, and the $x$ axis lists the models. The red line indicates the running time of the fastest model (132 sec).

#### Supplementary Tables

Table 1: Dataset of Bacteriophages from Extreme Environments. This table lists bacteriophages characterized by their genomic sequences available in the NCBI GenBank, along with their viral classification, habitats, NCBI accession numbers, and lifestyles.

| Viral family | Habitat | NCBI Acc. | Lifestyle |
|---|---|---|---|
| Autographiviridae | arsenic-rich env. (Psychrotolerant) | OM913597.1 | Virulent |
| | | OM913598.1 | Virulent |
| | | OM913599.1 | Virulent |
| unk. Caudoviricetes | Deep subseafloor sediments (381 m) | JF974314.1 | Temperate |
| | | JF974315.1 | Temperate |
| | Deep-sea (1000 m) | AB967974.1 | Temperate |
| Suviridae | Hadal, Mariana Trench (8,900 m) | OQ832096.1 | Temperate |
| Myoviridae | Biomining biotopes, highly acidic environments | JX507079.1 | Temperate |

#### References


[1] Balázs Ligeti, István Szepesi-Nagy, Babett Bodnár, Noémi Ligeti-Nagy, and János Juhász. ProkBERT family: genomic language models for microbiome applications. *Frontiers in Microbiology*, 14:1331233, 2024.

[2] Taku Kudo and John Richardson. Sentencepiece: A simple and language independent subword tokenizer and detokenizer for neural text processing. In *Proceedings of the 2018 Conference on Empirical Methods in Natural Language Processing: System Demonstrations*, pages 66–71, 2018.

[3] Rico Sennrich, Barry Haddow, and Alexandra Birch. Neural machine translation of rare words with subword units. *arXiv preprint arXiv:1609.08144*, 2016.

[4] Ofir Press, Noah A Smith, and Mike Lewis. Train short, test long: Attention with linear biases enables input length extrapolation. *arXiv preprint arXiv:2108.12409*, 2021.

[5] Tri Dao, Daniel Y Fu, Stefano Ermon, Atri Rudra, and Christopher Ré. Flashattention: Fast and memory-efficient exact attention with io-awareness. In *Proceedings of the IEEE/CVF Conference on Computer Vision and Pattern Recognition*, pages 12785–12795, 2022.

[6] Marie Lopez, Hugo Dalla-Torre, Liam Gonzalez, Javier Mendoza-Revilla, Nicolas Lopez Carranza, Adam Grzywaczewski, Francesco Oteri, Christian Dallago, Evan Trop, Hassan Sirelkhatim, et al. The Nucleotide Transformer: Building and Evaluating Robust Foundation Models for Human Genomics. 2023.

Table 2: Comprehensive Dataset Summary. This table provides a detailed summary of the datasets used in the study, categorized by the full sequence (contig) and segmented sequences (512bp and 1022bp). The datasets are split into training, validation, and test sets, with specific attention to temperate and virulent phages. The columns include the dataset name and split (training, validation, or test), the number of temperate and virulent sequences, and the total length of these sequences in megabases (Mb). This comprehensive summary highlights the diversity and scale of the datasets used for training, validating, and testing the phage lifestyle prediction models, illustrating the robustness and generalizability of the models across different sequence lengths and environmental contexts.

| **Dataset** | | **Size** | | **Length (Mb)** | |
|---|---|---|---|---|---|
| **Name** | **Split** | **Temperate** | **Virulent** | **Temperate** | **Virulent** |
| | | **Full Sequence - Contig** | | | |
| BACPHLIP | Train | 1112 | 756 | 56.74 | 64.86 |
| | Validation | 96 | 150 | 4.52 | 15.41 |
| *Escherichia* | Test | 200 | 194 | 8.58 | 13.82 |
| EXTREMOPHILE | Test | 10 | 6 | 0.48 | 0.26 |
| | | **Segments (512bp)** | | | |
| BACPHLIP | Train | 1098651 | 1259367 | 562.48 | 644.78 |
| | Validation | 43481 | 149995 | 22.26 | 76.8 |
| *Escherichia* | Test | 16766 | 26988 | 8.57 | 13.8 |
| EXTREMOPHILE | Test | 930 | 510 | 0.48 | 0.26 |
| | | **Segments (1022bp)** | | | |
| BACPHLIP | Train | 546118 | 626990 | 558.07 | 640.75 |
| | Validation | 21601 | 74983 | 22.07 | 76.62 |
| *Escherichia* | Test | 8404 | 13518 | 8.56 | 13.79 |
| EXTREMOPHILE | Test | 468 | 256 | 0.48 | 0.26 |
